# Hybrid peptide DNA nanomaterials enable potent and broad-spectrum virus neutralization

**DOI:** 10.1101/2025.07.21.666049

**Authors:** Saurabh Umrao, Abhisek Dwivedy, Payton L Haak, Dhanush Gandavadi, Laurie A Rund, Chi Chen, Lifeng Zhou, Jinwei Duan, Ying Fang, Andrew J Steelman, Xing Wang

## Abstract

The continued emergence of antigenic drift and drug-resistant viral strains highlights the need for antiviral strategies that deliver robust efficacy, broad subtype coverage, and minimal off-target toxicity. We demonstrate a potent and broad-spectrum strategy that employs hybrid biomaterials of Urumin (a host defense peptide) and a honeycomb (HC) DNA origami through spatially organized multivalent presentation for enhanced antiviral efficacy. Molecular dynamics simulations reveal that Urumin penetrates and destabilizes the hemagglutinin (HA) trimer core, disrupting influenza A viral (IAV) entry. Arranging Urumin in trimeric clusters on the HC enables potent multivalent binding to trimeric HAs on IAV, enhancing antiviral efficacy at nanomolar concentrations, ∼1,000-fold more effective than free Urumin. In vitro assays confirm HC-Urumin outperforms free Urumin in blocking viral entry and preserving cell viability in more IAV subtypes. In vivo studies show that compared to free Urumin, HC-Urumin treatment reduces disease severity, preserves physiological behavior, and decreases mortality in infected mice, while maintaining virus-specific adaptive immune responses without altering humoral immunity. Our study offers an advanced and effective materials platform and strategy for broad-spectrum, low-dose intervention against human and animal IAVs, which can be adapted to combat other viruses by patterning corresponding host defense peptides on custom designed DNA nanostructures.

## INTRODUCTION

Treatment and prevention of viral infections are of considerable importance due to their great potential to cause serious illnesses and transmit rapidly, leading to devastating impact on human/ food animal health and economy. For example, since introduced and eventually adapted to the human population, influenza A viruses (IAVs) have led to worldwide pandemics including the 1918 “Spanish flu” and the 2009 “Swine flu”, followed by their persistent circulation as seasonal influenza viruses [1-3]. Annually, seasonal IAV affects approximately one billion individuals worldwide, with 3–5 million cases progressing to severe illness and resulting in 290,000 to 650,000 respiratory-related fatalities (WHO) [4]. The recurring seasonal epidemics each year are a result of the continual evolution of seasonal IAV, which enables them to escape immunity generated from previous infections or vaccinations [5, 6]. During epidemics and pandemics, when vaccine production cannot keep pace with the outbreak, or there is a mismatch between predominant seasonal strains and vaccine formulations, anti-IAV treatments become the primary intervention. Current anti-IAV therapies target two viral proteins, M2 ion channels and neuraminidase (NA) [7, 8]. However, the emergence of escape mutations has rendered many viruses resistant to these drugs, reducing or negating their therapeutic efficacy [9].

To address these limitations, therapeutic strategies employing antiviral particles that target highly conserved functional domains of hemagglutinin (HA) protein involved in viral entry offer a promising alternative [10, 11]. IAV is well known to use its trimeric HA spike protein to attach to a host cell surface by recognizing sialic acid (SA)-terminated glycan receptors on the cellular membrane through multivalent binding mechanisms [12, 13]. Taking inspiration from such naturally occurring multivalency binding feature, a promising strategy to prevent viral infection involves the development of synthetic viral inhibitors that engage HA proteins in a multivalent format, thereby blocking their interaction with SA receptors and inhibiting the viral internalization. Various antiviral agents have been synthesized using scaffolds such as star-shaped polymers [14, 15], dendrimers [16, 17], and biocompatible nanoparticles [18-20] to selectively target the receptor-binding sites on the HA1 head region/domain. However, these scaffolds cannot precisely position the HA1-binding ligands to maximize their binding avidity. The HA1 head domain is subject to frequent mutations due to antigenic drift [21] and antigenic shift [22], often limiting the efficacy of immunity conferred by prior infections or vaccinations. Thus, the evolutionarily conserved HA2 stem domain, which undergoes much fewer mutations and is under less selective pressure, has emerged as a compelling target [23, 24]. An emerging approach for viral inhibitor development harnesses host defense peptides (HDPs), which are evolutionarily conserved components of innate immune systems across host species [25-31]. For example, Holthausen et al. identified a peptide, named Urumin from frog skin secretions, and demonstrated its potent neutralizing activity at micromolar concentrations against drug-resistant H1 influenza strains [32]. The study showed that Urumin exhibits inhibitory activity against H1 subtypes, but with minimal effects on H3 viruses. Although Urumin shows low cytotoxicity at therapeutic doses, its use at micromolar concentrations can promote non-specific interactions with host cell membranes, particularly in epithelial tissues, compromising membrane integrity. Prolonged or repeated exposure at such concentrations may approach cytotoxic thresholds, thereby narrow the therapeutic window and reduce the safety margin for clinical applications.

To address the suboptimal potency, limited subtype coverage, and off-target toxicity of the HDPs at therapeutic doses for viral neutralization, we turn our attention towards DNA nanomaterials, which offer a versatile platform for enabling pattern recognition-enabled multivalent interactions between the viral target and ligand [33]. We previously developed designer DNA nanostructures (DDNs) for the detection and neutralization of virus infections [34-38], and targeted drug delivery [39]. These DDNs leverage the principles of pattern-recognition enabled multivalent binding between spatially controlled molecular binders with viral antigens to achieve high binding avidity and specificity [40]. For example, DNA Net employs a grid-like trimeric architecture designed to pattern-match with the spatial arrangement of mobile spike proteins on the SARS-CoV-2, promoting cooperative binding for high-affinity interactions [36].

Building on the concept of pattern recognition-enabled multivalent interactions, we design a DNA nanomaterials-based, Urumin-enabled antiviral assay for advancing antiviral tools with broader virus neutralization by employing a honeycomb-shaped DDN (HC-DDN) framework [41].

Illustrated in **Fig. 1**, HC-DDN is engineered to display an array of trimeric binders, either the Urumin peptides [32] or fluorescent labelled HA-targeting DNA aptamers (called “V46”) reporters [42]. These ligands are precisely positioned at the center of each hexagonal unit of honeycomb, resulting in six uniformly distributed ligand trimeric clusters across the nanostructure, forming the HC-Urumin or HC-V46 constructs. Our data demonstrate that organizing Urumin peptides in a multivalent configuration on the DNA origami scaffold significantly enhances both potency and subtype coverage against IAVs, achieving over 98% inhibition of H1N1 and H3N2 subtypes at nanomolar concentrations while maintaining over 90% cell viability in murine and porcine respiratory epithelial models. In contrast, free Urumin peptide shows only ∼20% inhibition and ∼50% cell viability at micromolar doses. These findings highlight the synergistic benefit of precise nanoscale spatial arrangement and multivalency built on DNA nanomaterials platform, delivering up to a 1,000-fold improvement in antiviral efficacy compared to the monomeric peptide. Notably, the designer HC-Urumin platform enables broad-spectrum neutralization of both human and animal IAVs, overcoming the intrinsic limitations of free Urumin. The therapeutic promise of this approach is further supported by extensive molecular dynamics (MD) simulations and showcased in vivo using a murine H1N1 infection model, where treatment with HC–Urumin resulted in significantly reduced morbidity and mortality compared to free Urumin or untreated controls. Mice treated with HC–Urumin exhibited less weight loss and preserved burrowing behavior during the acute infection phase, reflecting a lower disease burden. Mortality was reduced to 20% in the HC–Urumin group, compared to 50% and 42% in the free Urumin and untreated groups, respectively, despite a high viral challenge dose (100 p.f.u.). Importantly, immunological profiling showed that HC–Urumin treatment did not impair virus-specific immune responses, as evidenced by comparable anti-HA IgG titers and robust CD8+ T cell responses in antigen recall assays. Collectively, these results indicate that HC–Urumin mitigates acute disease pathology without suppressing protective immunity, a critical consideration for platforms intended for prophylaxis or early therapeutic intervention.

**Figure 1.**
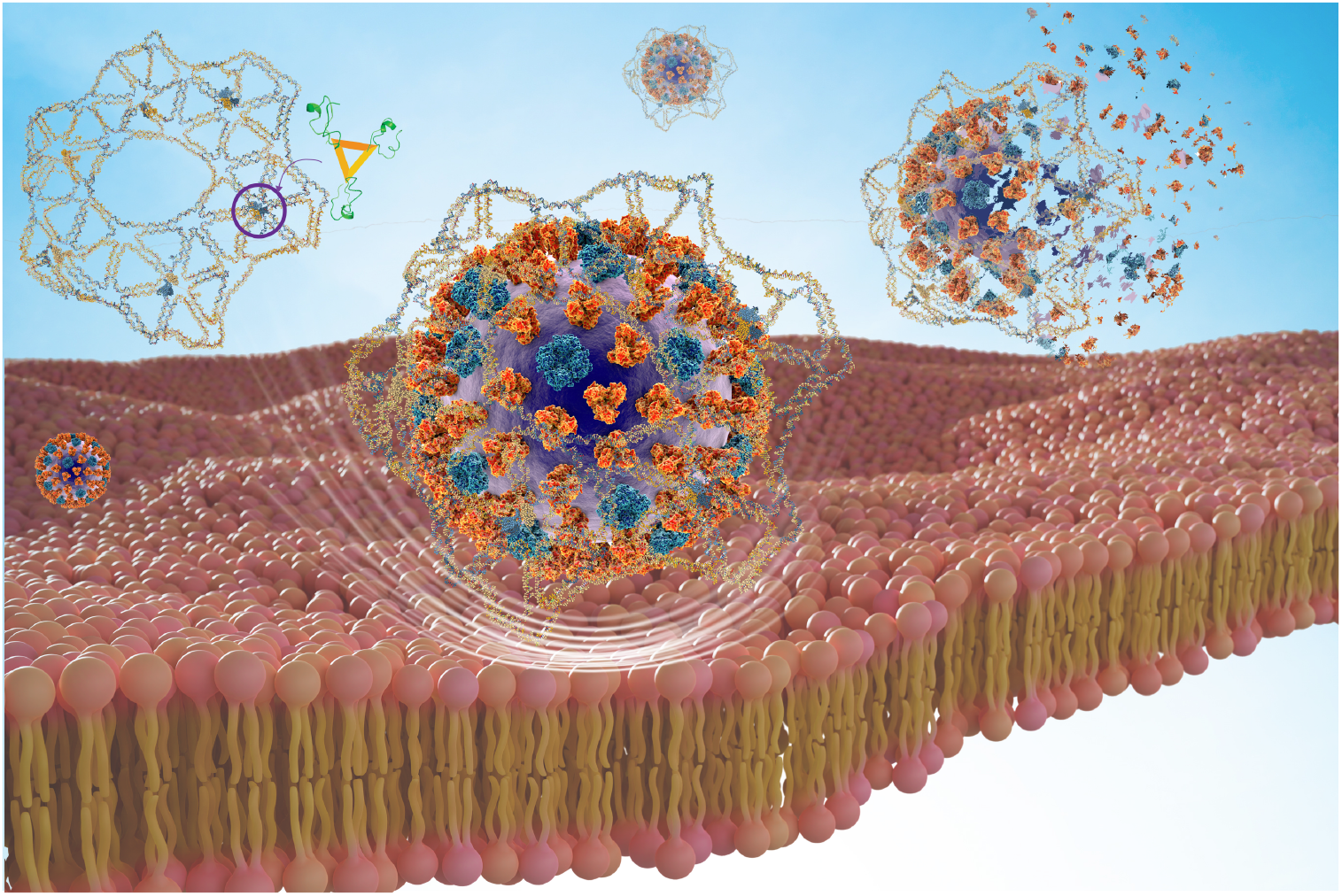
Honeycomb-shaped designer DNA nanostructure (HC-DDN) tethered with Urumin peptide enhances antiviral activity by blocking viral entry and significantly increasing virucidal activity.

In summary, this study presents a DNA nanomaterials–enabled antiviral strategy that markedly enhances the potency and breadth of biologically active peptide therapeutics through precise, multivalent pattern recognition. The programmable nature of the DNA nanomaterials-enabled platform technology permits facile adaptation to a wide variety of ligands, including alternative antiviral peptides, aptamers, and nanobodies, enabling rapid customization to diverse viral surface architectures. This plug-and-play strategy provides a versatile foundation for developing next-generation antivirals targeting respiratory pathogens such as respiratory syncytial virus (RSV) and coronaviruses.

## RESULTS AND DISCUSSION

### Computational analysis of molecular interactions between HA protein and Urumin peptide

To gain mechanistic insights into the antiviral action of the Urumin peptide against IAV of different subtypes, we employed a combination of molecular docking and molecular dynamics (MD) simulations to investigate its interactions with HA protein trimers across multiple viral subtypes. The HA protein trimers as ligands were obtained from the Protein Data Bank (PDB): H1N1 (PDB ID: 8SD4, A/Puerto Rico/8/1934)^54^ and H3N2 (PDB ID: 4WE9, A/Victoria/361/2011) [43]. The Urumin peptide was prepared as a receptor for docking, and the HADDOCK webserver [44] was used to generate protein-peptide complexes for both viral subtypes. Subsequently, all-atom molecular dynamics (MD) simulations [45] were performed for 192 ns (2,000 frames) to refine the H1N1–Urumin and H3N2–Urumin complexes and elucidate the molecular interactions and structural effects of Urumin on the HA protein trimers. Our analysis revealed that Urumin undergoes specific conformational changes, allowing it to penetrate the core of the HA trimer (**Movie S1**), potentially leading to the disruption of HA protein trimeric structure. Notably, Urumin predominantly interacts with the HA stem region, suggesting that its disruptive effects could extend to other viral subtypes, including H3N2 (**Movie S2**), thereby expanding its antiviral targets.

Analysis of Root Mean Square Deviation (RMSD) variation over the course of MD simulations for the HA amino acids suggests a destabilization of the Urumin-HA complex. However, Urumin-HA complex for H1N1 exhibits marginally higher deviations than for H3N2 **(Fig. 2a)**. This is also supported by an overall reduction in the number of H-bonds, suggesting the possible disruption in HA-trimeric state **(Fig. 2b)**. To support this hypothesis that the binding of Urumin and its penetration to the HA trimer could lead to the possible monomerization of HA, we analyzed the change in Solvent Accessible Surface Area (SASA). We observed increasing SASA values throughout the analysis, indicating the solvent interaction with the HA trimer’s core amino acids **(Fig. 2c)**. Furthermore, we analyzed the molecular surface of the Urumin at different frame numbers, 0, 1,000 and 1,999 of the MD simulation, and observed that Urumin not only penetrates towards the core of HA-trimer but also splits two adjacent HA helices **(Fig. 2d, Figs. S1-S2)**. However, this penetration was more pronounced in H1N1 HA. Molecular analysis of Urumin with HA at Frame 0 and 1,999 of the MD simulation reveals an increase in stronger interactions, such as ionic bonds and pi-interactions. This suggests that the Urumin not only enters the HA-trimer core but possibly keeps the core physically more open (in a so-called “closed” state that does not favor HA-cell interaction), subsequently increasing solvent accessibility, thus aiding to the disruption of the trimeric state **(Figs. S3-S4)**. We also observed key differences between the binding of Urumin to H1N1 and H3N2 HA trimer. While Urumin amino acids assume a flatter confirmation during the initial binding with H1N1 HA, for H3N2 HA, the amino acids form an arch, possible due to the difference in the nature of the interacting amino acids from HA **(Fig. S5)**. We hypothesize that this difference in the binding residues may affect the efficiency of Urumin in disrupting the HA trimeric state, thus in turn affecting its antiviral efficacy for different influenza subtypes.

**Figure 2.**
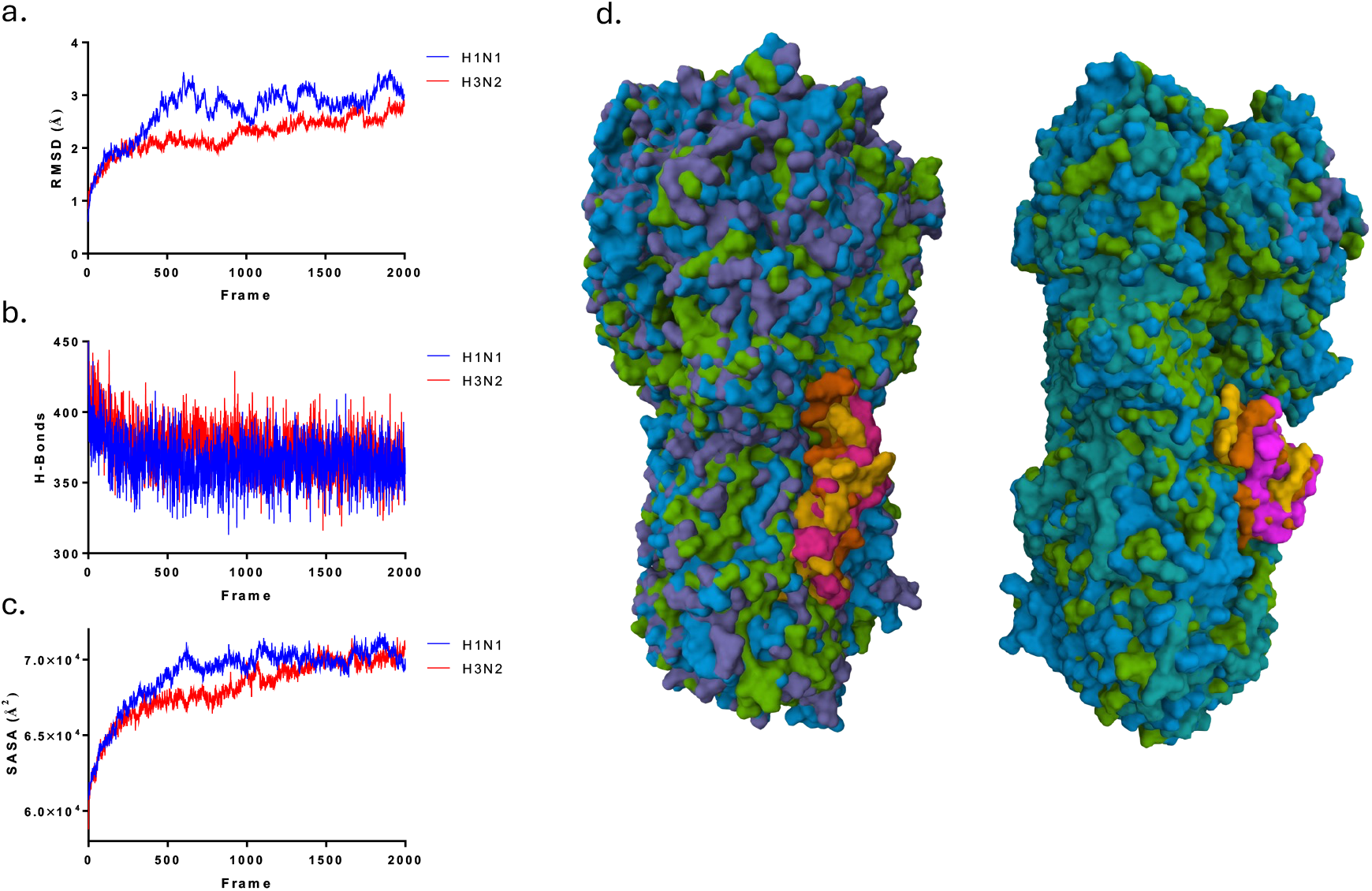
Molecular dynamics (MD) simulations suggest a potential mechanism of action for the Urumin peptide against influenza hemagglutinin (HA) trimers. (**a**) Root-mean-square deviation (RMSD), (**b**) number of hydrogen bonds (H-bonds), and (**c**) solvent-accessible surface area (SASA) analyses indicate structural destabilization of the hemagglutinin (HA) trimer complexes of both H1N1 and H3N2 influenza A subtypes following Urumin interaction. (**d**) Molecular surface representations illustrate the Urumin peptide interacting with the HA trimers of H1N1 (left panel) and H3N2 (right panel) at three time points during the MD simulation—initial frame 0 (pink), intermediate frame 1000 (yellow), and final frame 1999 (brown). The progressive penetration of the peptide into the HA trimer interface suggests a potential mechanism of trimer destabilization, which may contribute to the antiviral activity of Urumin.

### Design and simulation of the honeycomb-shaped designer DNA origami nanostructure

Following computational insights into the mechanism of Urumin interaction with the HA protein of different subtypes, we sought to improve this interaction through multivalent approach using a honeycomb-shaped designer DNA nanostructure (HC-DDN) as a modular scaffold. The goal was to spatially arrange multiple Urumin peptides on the HC-DDN platform to amplify binding avidity toward HA via multivalent interactions.

The HC-DDN construct comprises six hexagonal units, each with a minimum edge length of 42 base pairs, corresponding to approximately 14.28 nm. The initial design was generated using a wireframe double-crossover motif and a dual-graph scaffold routing strategy, which facilitates efficient scaffold routing within complex geometries. To improve structural flexibility and maintain high assembly fidelity, a discrete-edge design was employed, where each edge junction was modified to include seven thymidine bulges. This design choice, in contrast to continuous-edge architectures, provided localized bendability at crossover points for promoting the wrapping of virus particles while avoiding unpaired scaffold regions that could compromise structural stability.

Subsequent structural modifications and functionalization on HC-DDN platform were carried out using oxView2 software [46]. Single-stranded DNA linkers, complementary to those attached to antiviral peptides or signaling aptamers, were manually added to enable the site-specific incorporation of functional moieties. Specifically, we positioned either Urumin peptides or DNA aptamers (n = 18) in trimeric clusters at the center of each hexagonal unit. Each trimer exhibited an intra-cluster spacing of approximately 9.89 nm, with an inter-trimer spacing of ∼37.38 nm across the HC-DDN. This triangular mesh layout was designed to promote effective clustering of HA trimers on the curved viral envelope, thereby facilitating high-avidity interactions and enhancing antiviral efficacy.

To evaluate the structural integrity and dynamic behavior of the HC-DDN construct, we performed coarse-grained molecular dynamics simulations using the oxDNA framework [47]. Structural fluctuations were assessed through Root Mean Square Fluctuation (RMSF) analysis, while bond occupancy metrics were used to monitor base-pair stability over the simulation timeframe. RMSF values across the nanostructure ranged from 3.91 nm to 13.84 nm (**Fig. 3a**), with regions exhibiting minimal fluctuations represented in dark blue, and more mobile regions appearing in lighter blue to red. As anticipated, higher fluctuations were predominantly observed at terminal ends, where reduced crossover frequency leads to decreased structural rigidity. This pattern is consistent with the inherent flexibility of single-stranded DNA regions. To further assess base-pair integrity, we analyzed the bond occupancy parameter (**Fig. 3b**), which indicates how well the designed base-pair interactions are maintained under thermal fluctuations and molecular motion. Regions of persistent base-pairing (high occupancy) appear in red, while transient or unbound regions are depicted in white and blue, corresponding to partial and single-stranded segments, respectively. High occupancy indicates structural stability and correct folding of the HC-DDN. This is particularly important around functional attachment sites, where Urumin peptides or DNA aptamers are anchored, as maintaining local structural integrity is essential for spatial precision and preserving the biological activity of the HC-binder construct. Additionally, to visualize the system dynamics, we generated a simulation trajectory by centering the honeycomb structure within the simulation box and correcting for translational and rotational displacements (**Movie S3**). Collectively, these analyses confirm that the HC-DDN nanostructure retains its overall geometric fidelity and structural stability over the entire simulation period.

**Figure 3.**
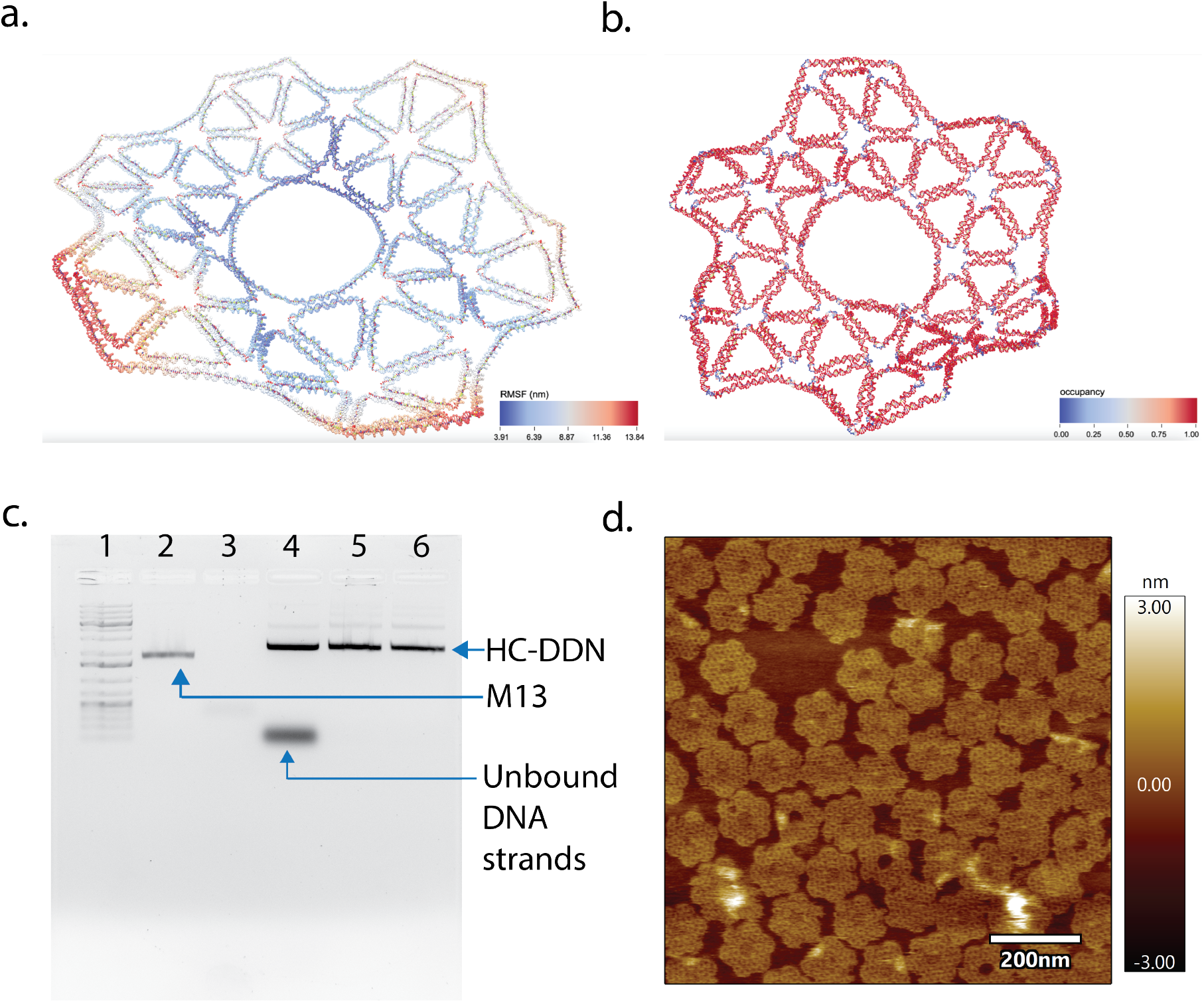
Design, simulation, and characterization of the honeycomb-shaped designer DNA nanostructure (HC-DDN). (**a**) Root Mean Square Fluctuation (RMSF) and (**b**) bond occupancy analyses were performed using OxDNA, a coarse-grained molecular dynamics (MD) simulation platform, over 10^9^ simulation steps with a 0.001 simulation integration time step. Dark blue regions indicate areas of minimal nucleotide fluctuation, while light blue regions represent segments with higher mobility, highlighting structural flexibility within the HC-DDN construct. (**c**) Experimental validation was performed using 1% agarose gel electrophoresis (AGE) to assess the formation and stability of the DNA nanostructure. Lane 1 contains the 1 kilobase-plus (1 kb+) DNA ladder; lane 2 shows M13 single-stranded DNA (ssDNA); lane 3 is a no-reagent blank control; lanes 4 and 5 contain HC-DDN samples before and after polyethylene glycol (PEG)-mediated purification, respectively; and lane 6 contains the HC-DDN conjugated with the Urumin peptide. (**d**) Atomic Force Microscopy (AFM) imaging was used to visualize the morphology and confirm the assembly of the HC-DDN nanostructure at nanoscale resolution.

### Synthesis and characterization of HC-DDN construct

To experimentally validate the successful assembly of the HC-DDN construct, we utilized 1% agarose gel electrophoresis (AGE) and atomic force microscopy (AFM) (**Figs. 3c–d**). Initial AGE analysis of the assembled product (lane 4) reveals the presence of excess staple strands. These were efficiently removed using a polyethylene glycol (PEG)-based purification protocol, as evidenced by the cleaner band observed in lane 5 of **Fig. 3c**. Following purification, the HC-DDN was conjugated with 18-fold copies of Urumin peptides to generate the multivalent HC-DDN– Urumin construct (lane 6), which was subsequently analyzed by AFM imaging (**Fig. 3d**). Both AGE and AFM imaging confirm the high-yield assembly of the HC-DDN construct. AFM imaging reveals well-defined, planar structures with smooth morphology and inherent flexibility, in agreement with the structural dynamics predicted by coarse-grained simulations. These findings support the successful synthesis of the HC-DDN scaffold and its suitability for downstream antiviral applications.

### In vitro evaluation of the HC–Urumin assay against murine H1N1 and H3N2 viruses

Flow cytometry has been utilized as a robust analytical technique for quantifying viral inhibition by assessing parameters such as viral load, infected cell populations, and drug efficacy [36, 38, 48, 49]. To enable precise monitoring of viral inhibition in conjunction with the HC–Urumin construct, we employed the V46 DNA aptamer, a FAM-labeled probe that specifically targets the highly conserved stem region of the HA protein [42]. The V46 aptamer functions both as a molecular binder and a fluorescent reporter to light up the virus particles. The V46 aptamer was conjugated to the HC-DDN scaffold using the same strategy applied for Urumin functionalization (**Fig. S6**), resulting in the HC-V46 construct, which was subsequently used as a reporter probe in viral infection assays.

We initiated the cell infection process by incubating murine-adapted IAVs with the HC–Urumin construct for 2 hours at 37°C, a temperature at which IAVs rapidly attach to host cells within 5 minutes and continue to enter cells for up to an hour [50]. This setup allowed us to compare the antiviral efficacy of free Urumin with that of the multivalent HC–Urumin construct in a multiplexed format. More specifically, cells were infected with H1N1 and H3N2 virions at a MOI of 100 treated with either free Urumin (2 μM and 4 μM, near the previously reported IC_50_ of 3.8 μM) or HC– Urumin (5 nM and 10 nM). Flow cytometry analysis was performed 1 hour post incubation reveals that HC–Urumin greatly outperformed free Urumin at both concentrations, effectively preventing infections across both viral subtypes (**Figs. 4a–b**). Notably, at 10 nM HC–Urumin, the mean relative fluorescence unit (∼30 RFU) was nearly indistinguishable from that of uninfected controls (**Figs. S7a–b**), suggesting near-complete viral neutralization (>98%) via multivalent engagement of HA trimers with Urumin peptides. In contrast, cells treated with free Urumin showed a median fluorescence intensity of ∼2000 RFU for both viral subtypes, indicating limited antiviral effect (∼20%), likely due to weaker, monovalent interactions. Importantly, HC–Urumin displays no preference between viral subtypes, exhibiting comparable inhibition of both H1N1 and H3N2 strains.

**Figure 4.**
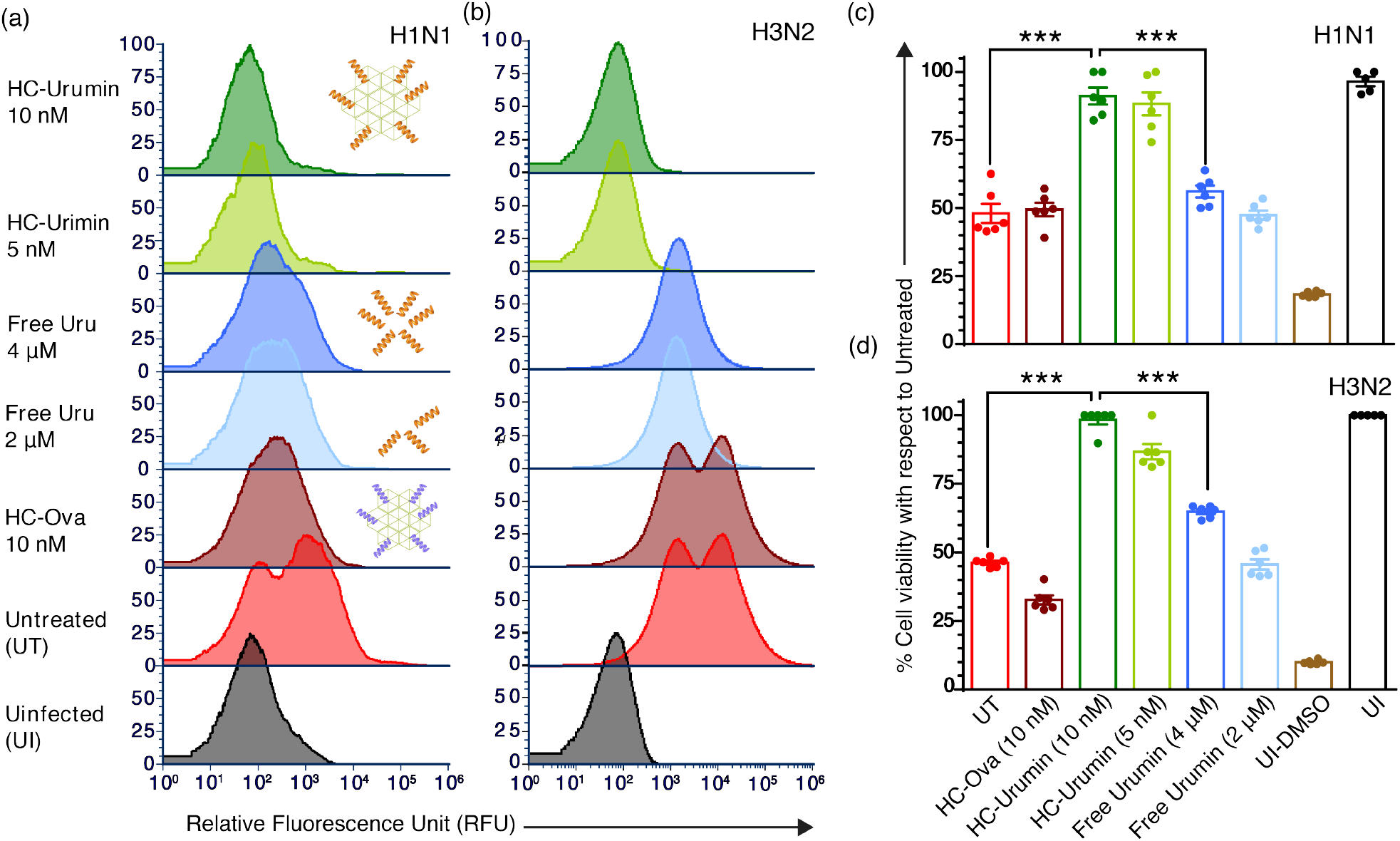
Honeycomb (HC)-shaped designer DNA nanostructure enhances the antiviral activity of Urumin and reduces viral entry of mouse-adapted influenza strains into host cells. (**a**) Flow cytometry analysis performed 1 h post-infection demonstrates enhanced antiviral efficacy of the HC-Urumin complex compared to free Urumin in preventing entry of mouse-adapted H1N1 influenza virions into host cells. HC construct complexed with a non-target peptide (ovalbumin or Ova), untreated virions, and uninfected cells were used as controls. (**b**) A comparable trend was observed with mouse-adapted H3N2 virions, although the enhancement in antiviral effect was less pronounced than that observed for H1N1. (**c, d**) Cell viability was assessed using the MTT assay 24 hours post-infection with H1N1 (**c**) and H3N2 (**d**) virions under various treatment conditions. The HC-Urumin construct significantly improved cell viability relative to free Urumin, suggesting reduced viral load and enhanced protective effect (DMSO treatment of uninfected cells serves as a positive control for cell death. Bars show mean and standard error, ^***^ denotes a p-value<0.0001).

To assess cytotoxicity, we performed an MTT assay comparing the effects of HC–Urumin, free Urumin, and HC–Ova (a non-HA targeting peptide used as a negative control) on cell viability. Cells infected with untreated or HC–Ova-treated H1N1 virus exhibited over 50% reduction in viability (**Fig. 4c**). In contrast, cells infected with H1N1 virions treated with 5 nM or 10 nM HC– Urumin maintained over 90% viability, comparable to uninfected controls. Free Urumin at 2 μM and 4 μM failed to offer similar protection, resulting in only 47.5% and 55% viability, respectively. Remarkably, HC–Urumin showed even greater efficacy against H3N2, restoring cell viability to 97.5% at 10 nM, dramatically outperforming free Urumin even at micromolar concentrations (**Fig. 4d**). These findings highlight the combinatorial effects of multivalent binding and spatial organization within HC-DDN structure, leading to enhanced, broader-spectrum antiviral activity and improved therapeutic efficacy exerted by Urumin peptide.

### Antiviral activity of the HC–Urumin assay against porcine H1N1 and H3N2 viruses

Different strains of IAV exhibit varying host specificity, which is largely governed by the affinity of the specific viral HA proteins to SA receptors on host cells [51]. Pigs, however, are uniquely susceptible to both avian and human IAVs [52], positioning them as critical intermediary hosts in the emergence of novel reassortant strains [53]. Additionally, the porcine immune system shares significant similarities with that of humans. Unlike mice, pigs naturally contract IAV and exhibit clinical symptoms comparable to those observed in humans [54]. As a result, pigs are widely recognized as a valuable model for studying human influenza pathogenesis and immune responses [55, 56]. These attributes underscore the relevance of evaluating antiviral candidates, such as HC–Urumin, against porcine-adapted IAV strains to better approximate therapeutic outcomes in humans.

Using the same approach as in murine-adapted IAV assays, we incubated porcine-adapted IAV (H1N1 and H3N2) with the HC–Urumin assay at 37 °C for 1 hour, followed by a 15-minute incubation with the HC– V46 reporter construct. Flow cytometry revealed that HC–Urumin (at both 5 nM and 10 nM concentrations) displays enhanced antiviral efficacy compared to free Urumin at micromolar concentration, significantly reducing infection in both porcine H1N1 and H3N2 subtypes (**Figs. 5a–b**). In H1N1-infected cells, HC–Urumin treatment reduces fluorescence intensity to ∼30 RFU, comparable to uninfected controls (**Fig. S8a**), whereas free Urumin at micromolar concentrations yields ∼150 RFU. Similarly, in H3N2-infected samples, HC–Urumin reduces the signal to ∼70 RFU compared to ∼150 RFU with free Urumin (**Fig. S8b**). These results demonstrate a marked enhancement in antiviral activity driven by the multivalent and spatially organized arrangement of Urumin on the HC-DDN scaffold. To further validate these findings, we assessed cell viability following infection with porcine-adapted H1N1 and H3N2. Cells infected with untreated virus or HC-Ova control exhibits ≥75% reduction in viability as determined by MTT assay (**Figs. 5c–d**). In contrast, cells exposed to virus treated with 10 nM HC-Urumin retains >95% viability for H1N1 and >80% for H3N2, levels comparable to uninfected controls. Free Urumin, in contrast, offers limited protection, with cell viability ranging from 50–55% for H1N1 and 45–60% for H3N2 across the 2–4 μM concentration range. Given the physiological similarities between pigs and humans, and the shared clinical manifestation of IAV infections, these findings suggest that HC-Urumin exhibits enhanced antiviral potency in more human-relevant systems. The observed broad-spectrum efficacy against both H1 and H3 subtypes further underscores its potential as a versatile and effective intervention strategy for influenza management.

**Figure 5.**
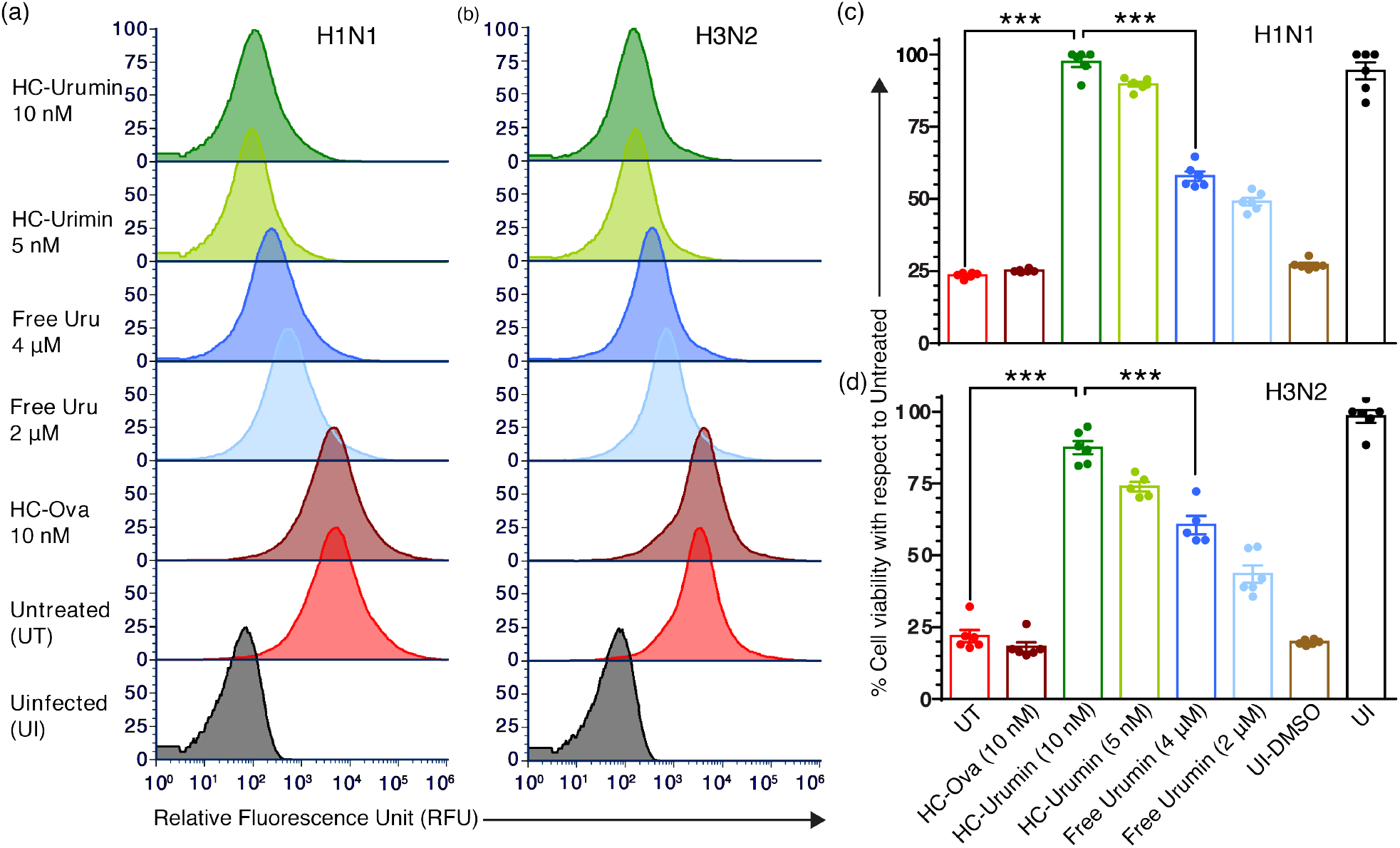
HC-DDN enhances the antiviral activity of Urumin and reduces swine-adapted influenza virus entry into host cells. (**a**) Flow cytometry analysis performed 1 h post-infection shows that Urumin complexed with HC-DDN exhibits enhanced antiviral efficacy against swine-adapted H1N1 virions compared to free Urumin. HC-DDN complexed with OVA peptide, untreated virions, and uninfected cells serve as controls. (**b**) A similar trend is observed for H3N2 virions, although the effect is less pronounced than that seen with H1N1. (**c, d**) MTT assays conducted 24 hours post-infection with H1N1 and H3N2 virions, respectively, demonstrate differential antiviral activity across treatment conditions. Urumin complexed with HC-DDN significantly improves cell viability, correlating with a reduced viral load compared to free Urumin. Importantly, HC-DDN–complexed Urumin maintains comparable antiviral efficacy even at half the concentration of previously reported therapeutic doses. Notably, this formulation shows greater antiviral potency against swine-adapted viruses than against mouse-adapted strains (see **Figure 4** for comparison) (DMSO treatment of uninfected cells serves as a positive control for cell death. Bars show mean and standard error, ^***^ denotes a p-value<0.0001).

### Viral treatment with HC–Urumin decreases disease severity in vivo

Given the promising in vitro results, we next sought to demonstrate the effects of HC-Urumin enabled treatment in vivo. For these experiments mouse-adapted human influenza virus (H1N1, PR8) was pre-incubated with either vehicle (PBS), free Urumin or HC–Urumin for two hours at 37°C then inoculated into mice by intranasal injection. For comparison, mice in control groups received either vehicle, free Urumin or HC–Urumin alone. Following inoculation, body weights and burrowing behavior were measured daily to assess disease outcomes [57, 58]. Mice in the control groups neither lost weight nor displayed signs of sickness (i.e. decreased burrowing behavior). However, mice inoculated with IAV characteristically exhibited substantial weight loss that was maximal at day 9 post infection (p.i.) and resolved by day 16 p.i. (**Fig. 6a**). The effects of IAV on weight loss was mitigated when the virus was pre-incubated with HC–Urumin, but not when it was pretreated with free Urumin (**Fig. 6a**). As observed with body weights, burrowing behavior decreased in mice inoculated with IAV, but nearly returned to baseline levels by day 15 p.i. Consistent with an ability for HC–Urumin to curb viral morbidity, mice in the HC–Urumin+IAV group burrowed significantly more than mice that received IAV alone (days 5-9 p.i.) and more than mice in the Urumin+IAV group (day 9 p.i.). Interestingly, mice in the Urumin+IAV group also burrowed significantly more than mice that received IAV alone on days 9-11 p.i. (**Fig. 6b**). Finally, over the course of the study approximately 42% (3/7) of mice inoculated with high dose IAV (100 p.f.u.) reached euthanasia criteria which represented a strong trend towards increased mortality over PBS inoculated control animals (p=0.06). Similarly, 50% (4/8) of mice inoculated with IAV pretreated with free Urumin ultimately reached euthanasia criteria, which represented a significant increase is mortality over mice control mice (**Fig. 6c**). Although not statistically significant, the incidence of moribund mice inoculated with IAV pretreated with HC–Urumin appeared reduced compared to other infected mice with only 20% (2/8) reaching euthanasia criteria. Collectively, these data demonstrate a protective effect on HC–Urumin on virus induced morbidity that superseded that of free Urumin alone.

**Figure 6.**
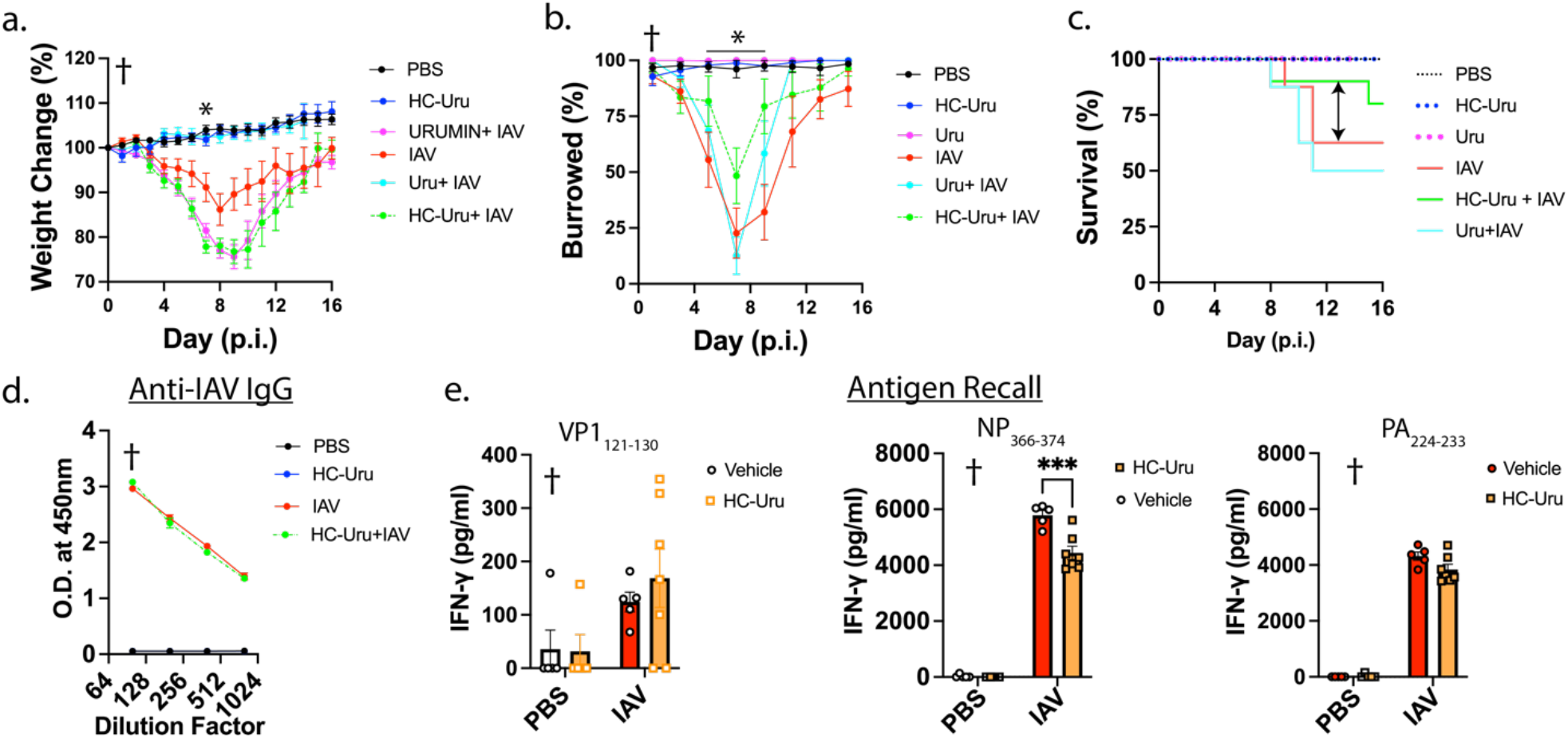
Viral pretreatment with HC-DDN-Urumin decreased disease severity in vivo. C57BL/6 mice were inoculated by i.n. route with sterile PBS, Urumin (180nM), HC-Urumin (10nM), IAV (PR8; 100 p.f.u.) or IAV pretreated with Urumin or HC-Urumin for 2h at 37°C. The effects of treatment and virus inoculation on percent change of body weight (**a**), burrowing behavior (**b**) and survival (**c**) is shown. Results are presented as means ± S.E.; n=4-8 per group. †, main effect of infection; *, p<0.05 HC-Urumin vs. both IAV and Urumin+IAV. The effect of HC-Urumin treatment on virus-specific HA IgG levels in serum (**d**) as determined by ELISA and (**e**) IFN-γ secretion following splenocyte stimulation with immunodominant MHC class I restricted peptide sequences specific to Theiler’s murine encephalomyelitis virus (VP2_121-130_) or IAV (H1N1 PR8; PA_224-233_ and NP_366-374_) for 72h at day 16 post infection (p.i.) is shown. Results are means ± S.E. n = 5–8 per group. †, main effect of infection; ^***^, p<0.001.

Finally, we sought to determine whether the treatment could affect the generation of virus-specific adaptive immunity. For these experiments, we focused on the effect of HC–Urumin, as it showed the most promising therapeutic efficacy. Regardless of treatment status, IAV inoculation robustly increased circulating levels of virus-specific IgG, which were not altered by treatment (**Fig. 6d**). Antigen recall assays performed on splenocyte cultures isolated from mice at day 16 p.i. indicated that splenocytes from infected mice produced low levels of IFN-γ following treatment with an irrelevant viral peptide (VP1_121-130_). In contrast, IFN-γ production was significantly increased in splenocytes from infected mice when stimulated with the IAV-specific immunodominant MHC class I restricted epitopes NP_366-374_ and PA_224-233_ and (**Fig. 6e**). Splentocytes from mice treated with HC–Urumin+IAV secreted less IFN-γ than those from IAV when stimulated with NP_366-374_ but not PA_224-233_ **(Fig. 6e**). Collectively, these data indicate that pretreatment with HC–Urumin did not affect the generation of a humoral response to the viral HA antigen while it remains unexplored if it impacts cytotoxic T cell responses.

## CONCLUSION

Urumin, a naturally occurring antiviral peptide derived from frog skin, targets a conserved region in the stem domain of the HA protein, disrupting viral integrity and preventing infection. While the monomeric form of Urumin effects against H1 subtypes at micromolar concentrations, its therapeutic application is limited by modest potency, subtype specificity, and cytotoxicity at higher doses. Elevated concentrations may also accelerate peptide degradation by proteases and increase the risk of non-specific binding to host proteins or cell surface factors, potentially reducing antiviral specificity and triggering unwanted immune responses. To address these challenges, we integrat Urumin with a designer DNA nanostructure platform, resulting in the HC– Urumin construct, which enables multiplexed presentation of Urumin in a spatially organized trimeric configuration. This architecture significantly enhances Urumin’s antiviral efficacy by increasing its local concentration and facilitating multivalent interactions with viral targets. The multivalent display allows for a reduced total peptide dosage while simultaneously amplifying its virucidal activity across two IAV subtypes, as supported by both computational and experimental evidence.

In vitro studies demonstrate robust inhibition of both murine- and porcine-adapted H1N1 and H3N2 IAV strains, with HC–Urumin outperforming free Urumin at significantly lower concentrations. Flow cytometry and MTT assays confirm reduced viral entry and improved cell viability, respectively, establishing the functional superiority of the multivalent platform compared to free Urumin. Importantly, in vivo evaluation in mice infected with H1N1 as an example reveals that pretreatment with HC–Urumin not only mitigates disease severity, reflected in improved body weight retention and burrowing behavior, but also reduces mortality trends. These protective effects were achieved without suppressing the generation of virus-specific humoral immunity and with only modest modulation of T cell responses, supporting HC–Urumin’s potential as an effective yet immunologically compatible antiviral strategy.

In conclusion, our study highlights how spatially organized multivalency enabled synthetic DNA nanostructure-based platform can be harnessed to enhance the therapeutic index of biologically derived antiviral agents. The HC–DDN scaffold offers a modular, tunable platform with broad adaptability to other peptide, aptamer, or nanobody payloads and to other pathogens, including those with pandemic potential. Future work will focus on optimizing pharmacokinetics, evaluating long-term immune memory and safety profiles in large animal models such as swine, and integrating this platform into diagnostic and therapeutic applications against a broader spectrum of respiratory viruses, including RSV and coronaviruses.

## ACKNOWLEDGEMENTS

The authors would like to thank Jayleen Ortiz for her excellent technical support. The study was supported in part by NIH (R21AI166898) and ACES FIRE Grant (no. ILLU-538-932).

## Notes

### Competing Interest Statement

The authors have declared no competing interest.

